# Influence of different pharmaceuticals on the germination and early development of two leafy vegetable species

**DOI:** 10.64898/2026.04.08.717353

**Authors:** Lukas Brokate, Caroline Müller

**Affiliations:** Bielefeld University, Universitätsstr. 25, 33615 Bielefeld; Joint Institute for Individualisation in a Changing Environment (JICE), University of Münster and Bielefeld University, Bielefeld, Germany

**Keywords:** Pharmaceuticals, Phytotoxicity, Seed germination, Root elongation, Food crops, Leaf vegetables, Early development

## Abstract

Pharmaceuticals are becoming increasingly prevalent in the environment, yet their effects on terrestrial plants, especially during early development, are poorly investigated. In this context, leafy vegetables are of particular interest because they tend to accumulate more pharmaceuticals than other crops. This study investigated the impacts of six pharmaceuticals of different classes commonly detected in soils and water on seed germination and early seedling growth of the leafy vegetables bok choy (*Brassica rapa* subsp. *chinensis*) and spinach (*Spinacia oleracea*) under controlled conditions. Seeds were exposed to different concentrations of the non-steroidal anti-inflammatory drugs (NSAIDs) ibuprofen, naproxen, diclofenac, or salicylic acid, the antiepileptic drug carbamazepine, or the antibiotic ciprofloxacin, and germination rates, root and shoot lengths, biomass allocation, cotyledon development, and lateral root formation (in bok choy only) measured after seven days. While germination was unaffected, early development parameters showed species-specific responses. In bok choy, high concentrations of NSAIDs and ciprofloxacin led to an increased shoot biomass and cotyledon area but a reduced primary root growth and lateral root formation, while carbamazepine had no effect. The contrasting effects on aboveground versus belowground organs of different pharmaceuticals suggest an interference with hormonal regulation, especially auxin. Spinach showed less responses than bok choy, with root length being rather increased by some NSAIDs. These results indicate that sensitivity to pharmaceuticals begins after germination and depends on both the chemical properties of the compound and the plant species. The study highlights the value of systematic comparative testing of pharmaceuticals across plant species.

## 1. Introduction

Pharmaceuticals are widely used in modern society for the prevention and treatment of human and animal diseases and have become indispensable for modern healthcare. Estimates from 2014 suggest that around 4,000 active pharmaceutical ingredients (APIs) were used for human and veterinary applications worldwide, with an annual production of approximately 100,000 tons (Weber et al., 2014). More recent data indicate that already in Germany, around 3,000 APIs were in use in 2025, with an annual consumption exceeding 35,000 tons (German Environment Agency, 2025). Beyond their intended therapeutic use, pharmaceuticals have been considered as emerging environmental pollutants for several decades (Richardson and Bowron, 1985). Based on environmental monitoring data from 89 countries collected in the PHARMS-UBA database, a total of 992 different APIs and transformation products have been detected worldwide across 61 environmental matrices, including soils (Graumnitz and Jungmann, 2021). Despite extensive data on their environmental occurrence, little is known about their impacts on organisms that grow in contaminated soils, such as (crop) plants.

Pharmaceuticals enter the environment through multiple ways, including through wastewater from improper use, disposal, and human excretion in private households, but also in large quantities via industrial and hospital effluents (Weber et al., 2014; Gworek et al., 2021). Wastewater treatment plants are unable to completely remove most pharmaceutical compounds, with average removal efficiencies of ∼50% for antibiotics and approximately 30 - 40% for analgesics, anti-inflammatory drugs, and β-blockers (Deblonde et al., 2011). Worldwide, the majority of pharmaceuticals in the environment can be detected in liquid compartments (Graumnitz and Jungmann, 2021), which may explain why earlier studies focused primarily on impacts of pharmaceutical contaminants in aquatic systems (Kümmerer, 2010; Sleight et al., 2023). In contrast, ca. 30% are detected in solid environmental compartments, with around 60% of measured concentrations in soil samples exceeding the detection limit (Graumnitz and Jungmann, 2021). Research on the occurrence of pharmaceuticals and effects in terrestrial environments, including impacts on crops, only started in the 2000s, and their impacts remain poorly understood (Sleight et al., 2023).

Pharmaceuticals reach crops through different pathways, such as groundwater and irrigation as well as agricultural practices, particularly the application of manure and sewage sludge (Chia et al., 2025; Amin and Rahman, 2026). Despite major differences in environmental persistence and degradability, pharmaceuticals remain permanently in soils due to continuous inputs, leading to pseudo-persistence (Garduño-Jiménez and Carter, 2024). This continuous presence is of relevance, as numerous studies have demonstrated that pharmaceuticals present in soils can be taken up and accumulated by plants (Keerthanan et al., 2021). Once taken up, they can affect key plant traits such as photosynthetic performance, oxidative stress responses, growth, energy metabolism, and hormonal regulation (Garduño-Jiménez and Carter, 2024). However, these impacts highly depend on the class of pharmaceuticals due to their different modes of action and target structures in the human body and potentially in other organisms (Garduño-Jiménez and Carter, 2024). For example, non-steroidal anti-inflammatory drugs (NSAIDs), such as ibuprofen, disrupt auxin transport and actin dynamics in plants (Tan et al., 2020). In contrast, antibiotics may harm plants by decreasing the synthesis of chlorophyll, as well as by impacting their mutualistic microbiota (Carballo et al., 2022). Moreover, the responses vary between plant species and due to crop characteristics (Sunyer-Caldú et al., 2023; Kummerová et al., 2024). For instance, species of Brassicaceae have been shown to be more sensitive to antibiotics than members of Asteraceae, particularly regarding elongation growth (Carballo et al., 2022). In comparison, little is known about the effects of pharmaceuticals on seed germination and early plant development, especially in leafy vegetables, although they tend to take up and accumulate more pharmaceuticals than other crop types (Christou et al., 2019). Comparative studies are needed, in which different pharmaceuticals are tested in equal concentrations on different plant species under similar conditions.

In the present study, we tested thus the impacts of six pharmaceuticals with partly different mode of action on seed germination and early development of two leafy vegetable crop species, bok choy (*Brassica rapa* subsp. *chinensis*, Brassicaceae) and spinach (*Spinacia oleracea*, Amaranthaceae). Previous studies have demonstrated that these two species can take up and translocate pharmaceuticals, such as carbamazepine, to the leaves (Li et al., 2018; Kodešová et al., 2019). The pharmaceuticals used in this study were selected mainly based on frequent detection in the environment, their potential environmental risks, and insights from publications on potential impacts on plants. Therefore, the four NSAIDs ibuprofen, naproxen, diclofenac, and salicylic acid, the antiepileptic drug carbamazepine, and the antibiotic ciprofloxacin were chosen. Ibuprofen, naproxen, diclofenac, and carbamazepine have been reported as four of the five most frequently detected pharmaceuticals in surface, groundwater, and drinking water worldwide (Graumnitz and Jungmann, 2021). Recent studies on diclofenac and naproxen (10 mg/L) found no effect on seed germination of several monocot and dicot species (Kummerová et al., 2024), but a decrease in root length and biomass was observed in several-day-old *Solanum lycopersicum* (2 mg/L) (Siemieniuk et al., 2021). Furthermore, ibuprofen (40 µM) caused significant suppression of lateral root formation in *Arabidopsis thaliana* seedlings (Tan et al., 2020). Hardly any research exists on the effects of carbamazepine and ciprofloxacin on seed germination or early development.

We studied the impacts of the six pharmaceuticals in four concentrations. Based on previous findings in other systems, we hypothesized that none of the six pharmaceuticals influences the germination rate, but that they negatively affect the development of both plant species, especially root elongation, that the magnitude of these effects depends on the (class of) pharmaceutical and its concentration, and that effects are plant species-specific.

## 2. Materials and methods

### 2.1. Seeds

Bok choy ‘Joi Choi F1’ and spinach ‘Apollo F1’ seeds supplied by FLOVEG GmbH (Kall, Germany) were used in this study. Seeds were stored dry at room temperature. According to the supplier’s specifications and EU seed standards, the certified germination rate was 90% for bok choy ‘Joi Choi F1’ and 85% for spinach ‘Apollo F1’.

### 2.2. Pharmaceuticals and solutions

Ibuprofen sodium salt, naproxen sodium salt, diclofenac sodium salt, and sodium salicylate were obtained from Sigma-Aldrich (St. Louis, MO, USA). Sodium salts were used due to their better solubility. Sodium salicylate was used instead of acetylsalicylic acid because acetylsalicylic acid is rapidly metabolized into salicylic acid in the human body, leading to higher environmental concentrations of salicylic acid than acetylsalicylic acid (Nunes et al., 2015; Zivna et al., 2016). Ciprofloxacin, carbamazepine, and 2,3,5-triiodobenzoic acid (TIBA) were purchased from Thermo Fisher Scientific (Waltham, MA, USA). TIBA served as a positive control, as this acid is known to have highly negative effects on root elongation (Tan et al., 2020). All compounds had a purity ≥ 98 %.

Stock solutions were prepared by dissolving 5 mg of each pharmaceutical in 500 mL of Millipore water, resulting in a concentration of 10 mg/L. Test solutions of 0.01, 0.1, 1, and 10 mg/L were prepared by serial dilution. These concentrations were chosen based on reported maximum environmental concentrations of the investigated pharmaceuticals and extended to higher levels to investigate potential dose-dependent effects (Xie et al., 2012; Ashfaq et al., 2017). Millipore water was used as the negative control, while TIBA as positive control was prepared in a 10 mg/L solution. Carbamazepine, ciprofloxacin, and TIBA solutions were sonicated twice for 5 min to ensure complete solution. The pH values were measured for all solutions and were comparable among serial dilutions within each pharmaceutical (Table S1).

### 2.3. Germination assay

To achieve a reliable germination, spinach seeds required additional pretreatment steps and different growth parameters compared to bok choy, based on preliminary experiments. Spinach seeds were pretreated by storage in darkness at 4 °C for 7 days before use and were subsequently soaked for 24 h in the respective pharmaceutical solutions. Bok choy seeds were used without prior pretreatment. Seeds of both species were surface sterilized with 1% sodium hypochlorite for 10 min, rinsed 3 times with distilled water for 1 min each, and air dried before further use.

Germination assays were performed in Petri dishes (94 mm diameter × 16 mm height, Greiner Bio-One, Kremsmünster, Austria) with two filter papers (85 mm diameter; Whatman Grade1, Cytiva, Marlborough, MA, USA) for bok choy and one for spinach. Ten seeds were placed in each Petri dish and 5 mL of a given solution for bok choy or 0.5 mL for spinach added. Each pharmaceutical was tested at each concentration with eight replicates per plant species as well as eight replicates for each negative and positive control. Pharmaceuticals were tested sequentially. After use, each Petri dish was cleaned with distilled water and 70% ethanol and used in the next assay with the same concentration with a different pharmaceutical. For each species, Petri dishes were randomly arranged in two climate cabinets (CLF PlantClimatics GmbH, Wertingen, Germany; one cabinet per species) under dark conditions at 80% relative humidity. Temperatures were set to 20 °C for bok choy and 16 °C for spinach. After 3 days, light was switched on for bok choy for 16 h per day and 2 mL of the respective solution were added to the Petri dishes with spinach seeds. On day 6, light was also switched on for spinach for 16 h per day. Photosynthetically active radiation (PAR) values ranged between 207.2 and 257.2 µmol/m² * s in both climate cabinets. Germination rate was determined every 24 h. Seeds were considered germinated when the radicle was longer than 2.0 mm.

After 7 days, five fully developed germinated seeds from each Petri dish were randomly chosen and directly dissected into roots and shoots by cutting at the root–shoot junction. Fully developed seedlings were defined as individuals in which roots, hypocotyl, and cotyledons were clearly distinguishable, and the cotyledons were green and visibly developed. The plant parts were gently dried on paper and then weighed using an analytical balance with a precision of 0.1 mg (CPA224S-0CE, Sartorius AG, Göttingen, Germany) to determine the fresh biomass. Based on these data, the ratio of the root to the shoot biomass was calculated. To measure root and hypocotyl lengths in both species, cotyledon length in spinach, and cotyledon area in bok choy, seedlings were placed on a flat surface, covered with a glass plate, and photographed at a fixed distance (Fig. 1). A scale in the form of a millimeter grid was included in each image. Lengths and areas were measured using ImageJ software (version 1.54g; National Institutes of Health, Bethesda, MD, USA). For hypocotyl length of spinach, the longest hypocotyl per seedling was considered. For the analysis of the bok choy cotyledon area, identical ImageJ settings were applied to all images to ensure consistent detection and comparability across samples.

**Fig. 1.**
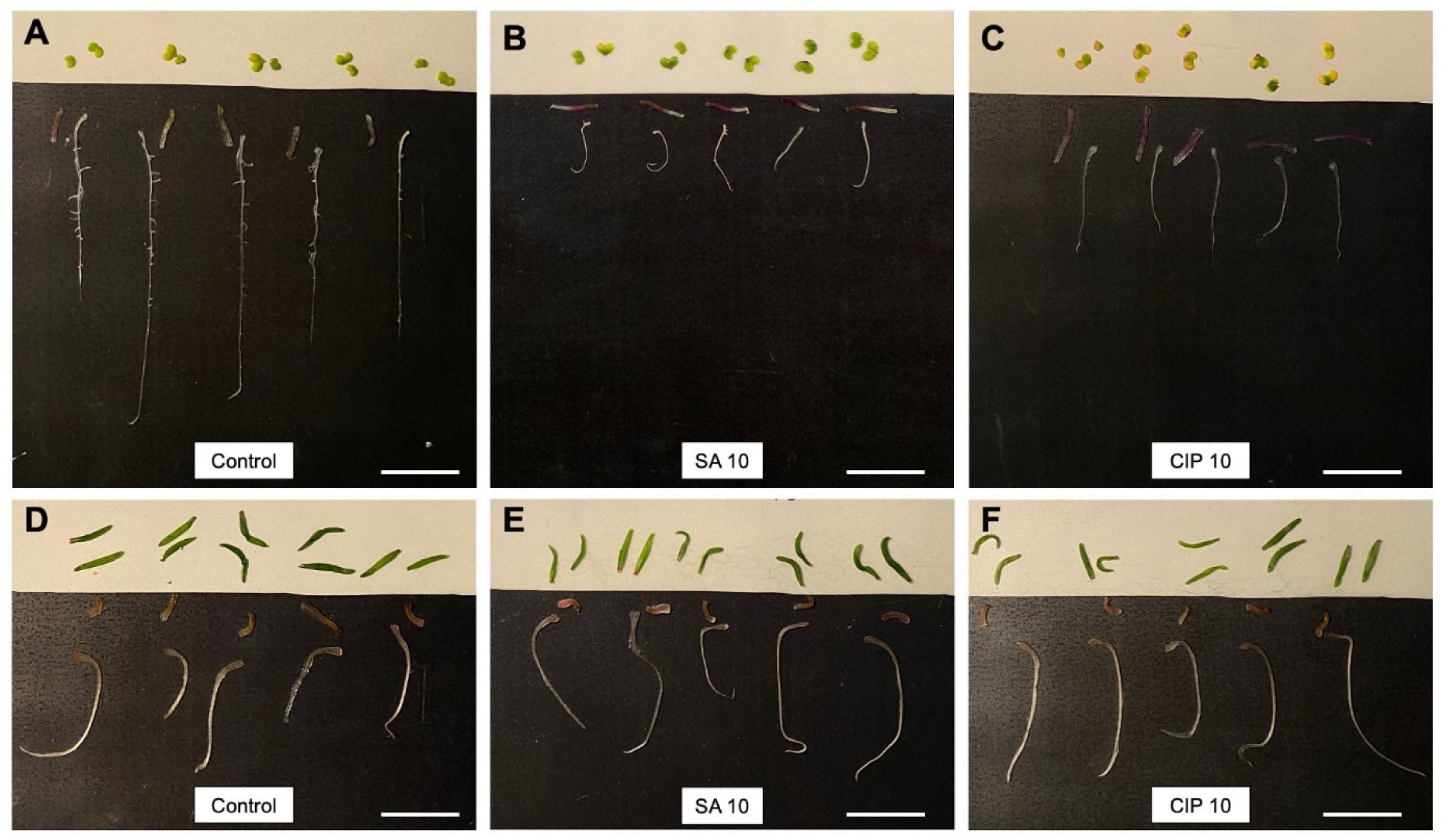
Visual analysis of dissected seedlings after exposure to pharmaceuticals. Seven-day-old seedlings of *Brassica rapa* subsp. *chinensis* (bok choy, A–C) and *Spinacia oleracea* (spinach, D–F) were dissected into roots, hypocotyls, and cotyledons after exposure to water (control), 10 mg/L salicylic acid (SA 10), or 10 mg/L ciprofloxacin (CIP 10) solutions. These pictures were used to measure lengths and areas of the plant parts with ImageJ software, using different backgrounds for better visibility. The scale bar is 2 cm.

### 2.4. Lateral roots assay

To test the impacts of the pharmaceuticals on the development of lateral roots, a separate assay was performed using bok choy only, as spinach did not develop any visible lateral roots under the experimental conditions. All experimental conditions were identical to those used in the germination assay for bok choi, but with three replicates per pharmaceutical and concentration and all tests running in parallel. After 7 days, the lateral roots of each germinated seed were counted and classified into the three categories C1 (< 1 mm), C2 (1 - 5 mm), and C3 (> 5 mm), using a millimeter grid.

### 2.5. Statistical analyses

All statistical analyses and data visualization were performed in R (version 4.4.3; R Core Team, 2025). The packages multcompView (Graves et al., 2024), rstatix (Kassambara, 2025), tidyverse (Wickham, 2023), and readxl (Wickham and Bryan, 2025) were used. For each pharmaceutical and species separately, differences among concentrations were evaluated using Kruskal–Wallis tests, followed by Dunn’s post hoc tests with Bonferroni-Holm correction (α = 0.05).

## 3. Results

### 3.1. Impacts of pharmaceuticals on germination

Cumulative germination curves were largely overlapping among the control, the different concentrations of all pharmaceuticals, and TIBA for both species (Fig. 2). For bok choy, more than 80% of the seeds germinated within the first 24 h in all treatments, except in the 1 mg/L ciprofloxacin treatment, where somewhat less germinated. By 48 h, germination exceeded 97% across all treatments and concentrations, including the control and TIBA. Standard deviations were low overall, with a higher variability primarily observed after 24 h, especially in the salicylic acid, carbamazepine, and ciprofloxacin treatments (Fig. 2).

**Fig. 2.**
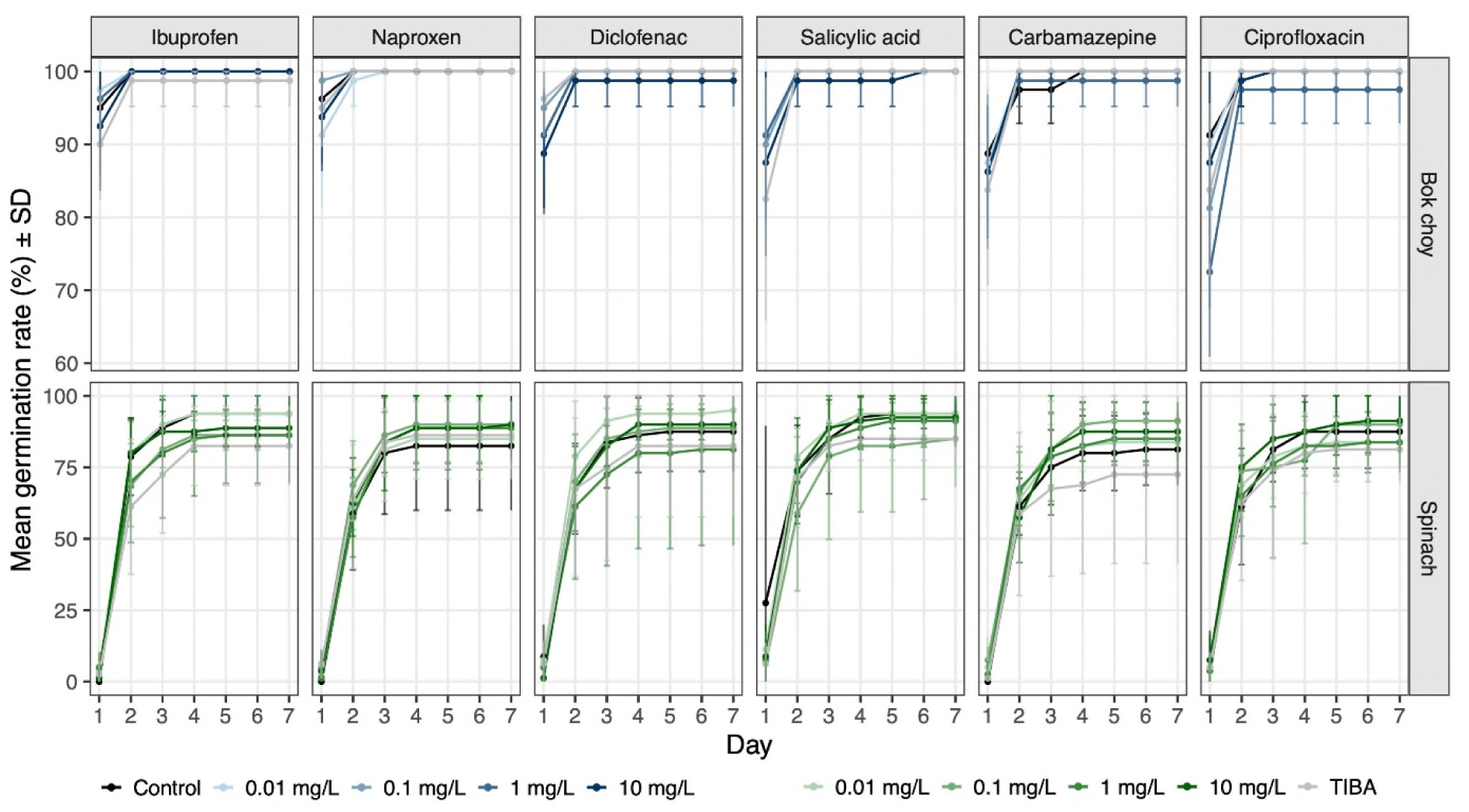
Mean cumulative germination rate (%) of *Brassica rapa* subsp. *chinensis* (bok choy) and *Spinacia oleracea* (spinach) over seven days when exposed to six pharmaceuticals. Pharmaceuticals were tested in four concentrations, using water as negative control, and 10 mg/L 2,3,5-triiodobenzoid acid (TIBA) as a positive control, with *n* = 8 Petri dishes, with 10 seeds per dish, per solution. Data are shown as line plots with points representing mean ± SD.

For spinach, germination onset occurred slightly later and proceeded more gradually than for bok choy. Most seeds germinated between 24 h and 48 h, whereby many seeds still germinated between 48 h and 72 h. After 96 h almost no additional germination was observed and cumulative germination rates ranged from 80 to 95% across all treatments, except for the TIBA treatment within the carbamazepine series, where germination was lower. Slight differences in germination between concentrations within pharmaceuticals were observed, without any concentration-dependent pattern. Standard deviations were higher than those observed for bok choy (Fig. 2).

### 3.2. Impacts of pharmaceuticals on early development of bok choy seedlings

Exposure to ibuprofen, naproxen, diclofenac, salicylic acid, and ciprofloxacin affected early developmental parameters in bok choy seedlings in different ways, with significant effects mainly occurring at higher concentrations (1 and/or 10 mg/L). Seedling development at lower concentrations of 0.01 and 0.1 mg/L, as well as under all carbamazepine treatments, did not differ significantly from the negative controls (Fig. 3A-C).

**Fig. 3.**
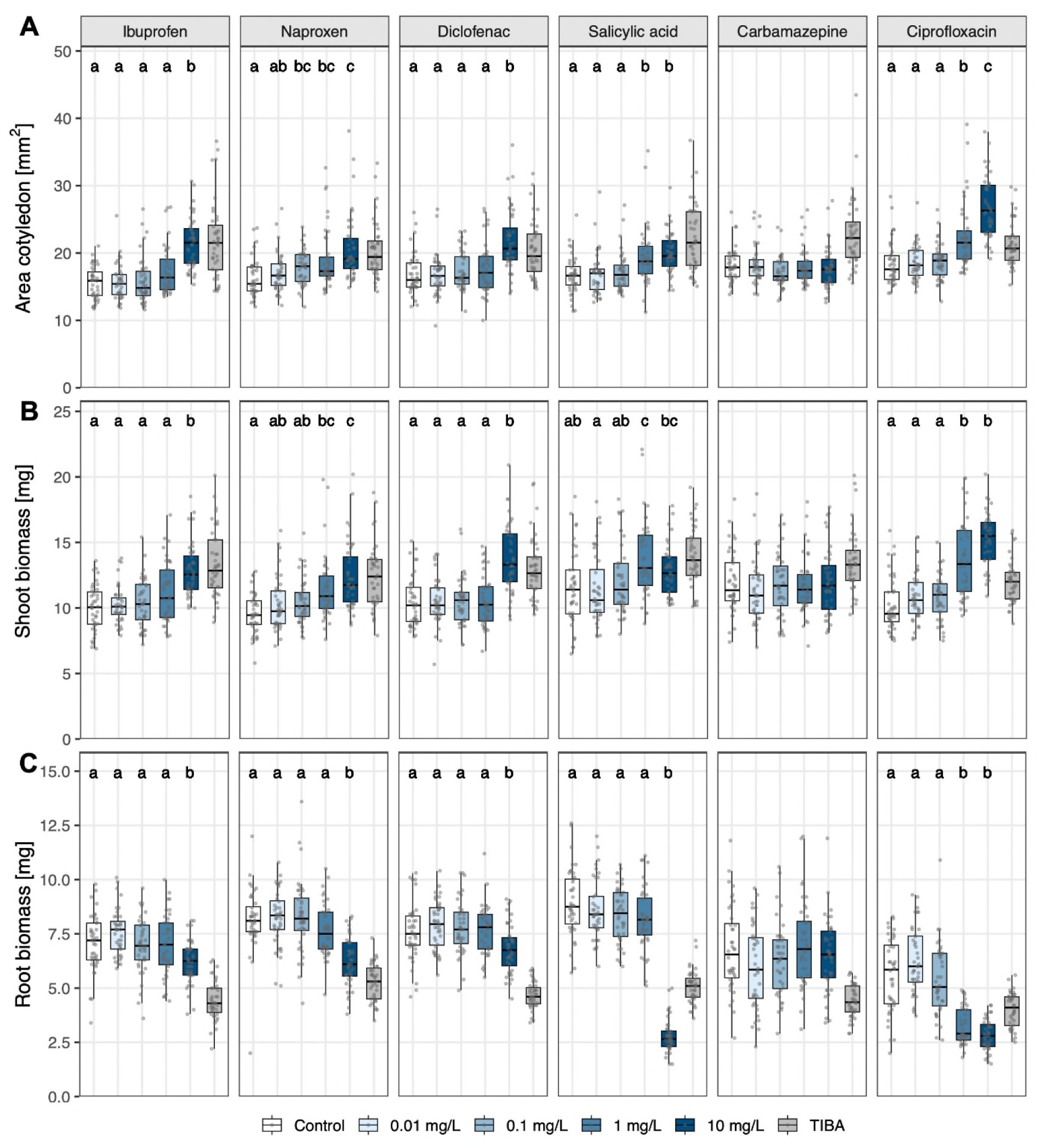
Area of cotyledons (A), root biomass (B) and shoot biomass (C) of seven-day-old seedlings of *Brassica rapa* subsp. *chinensis* (bok choy) exposed to six pharmaceuticals. Pharmaceuticals were tested in four concentrations, using water as negative control, and 10 mg/L 2,3,5-triiodobenzoid acid (TIBA) as a positive control. Data are presented as boxplots, with medians, interquartile ranges (IQR, boxes), and whiskers extending to the most extreme data points within 1.5×IQR. Individual values are plotted as dots; *n* = 40 per concentration (from 8 Petri dishes with 10 seeds per dish, 5 germinated seedlings were randomly taken per dish). Different letters indicate statistically significant differences within each pharmaceutical based on Dunn’s post hoc tests following a Kruskal–Wallis test (Bonferroni-Holm-adjusted *p* < 0.05).

The area of cotyledons (Fig. 3A) and the shoot biomass (Fig. 3B), as well as the length of the hypocotyls (Fig. S1) were enhanced across all tested pharmaceuticals, except carbamazepine, relative to the control. When exposed to ciprofloxacin at 10 mg/L, differences were particularly high, with a median shoot biomass of the bok choy seedlings being more than 60% higher than in the control, while the median cotyledon area was ca. 50% higher compared to the control. In addition, the cotyledons of seedlings exposed to this treatment appeared yellowish (Figs 1C; 3A-B).

In contrast, the root biomass (Fig. 3C) as well as the root length (Fig. S2) were lower at the highest tested concentration (10 mg/L) of all pharmaceuticals except carbamazepine compared to the negative controls. In addition, when exposed to 1 mg/L of ciprofloxacin, root biomass was also already ca. 50% lower than in the control. Bok choy seedlings exposed to 10 mg/L salicylic acid showed the highest difference in root biomass compared to the control, with a median root biomass of just 30% of that of the control (Figs 1B; 3C).

### 3.3. Impacts of pharmaceuticals on early development of spinach

Treatment of spinach seeds with the different pharmaceuticals generally had only minor effects on early developmental parameters (Fig. 1D-F), such as the root length (Fig. 4A) and the root-to-shoot biomass (Fig. 4B). Minor effects were also observed for the cotyledon length, with a significant difference detected only for a single pairwise comparison of two treatments with different naproxen concentrations (Fig. S3). Under exposure to ibuprofen and naproxen, the root length was significantly increased at 10 mg/L compared to the negative control, whereas under exposure to diclofenac a significant increase was observed just at 0.1 mg/L. In contrast, no significant differences among concentrations were detected in root length of spinach seedlings exposed to salicylic acid, carbamazepine, or ciprofloxacin, while the TIBA exposure mainly resulted in a decreased root length (Fig. 4A).

**Fig. 4.**
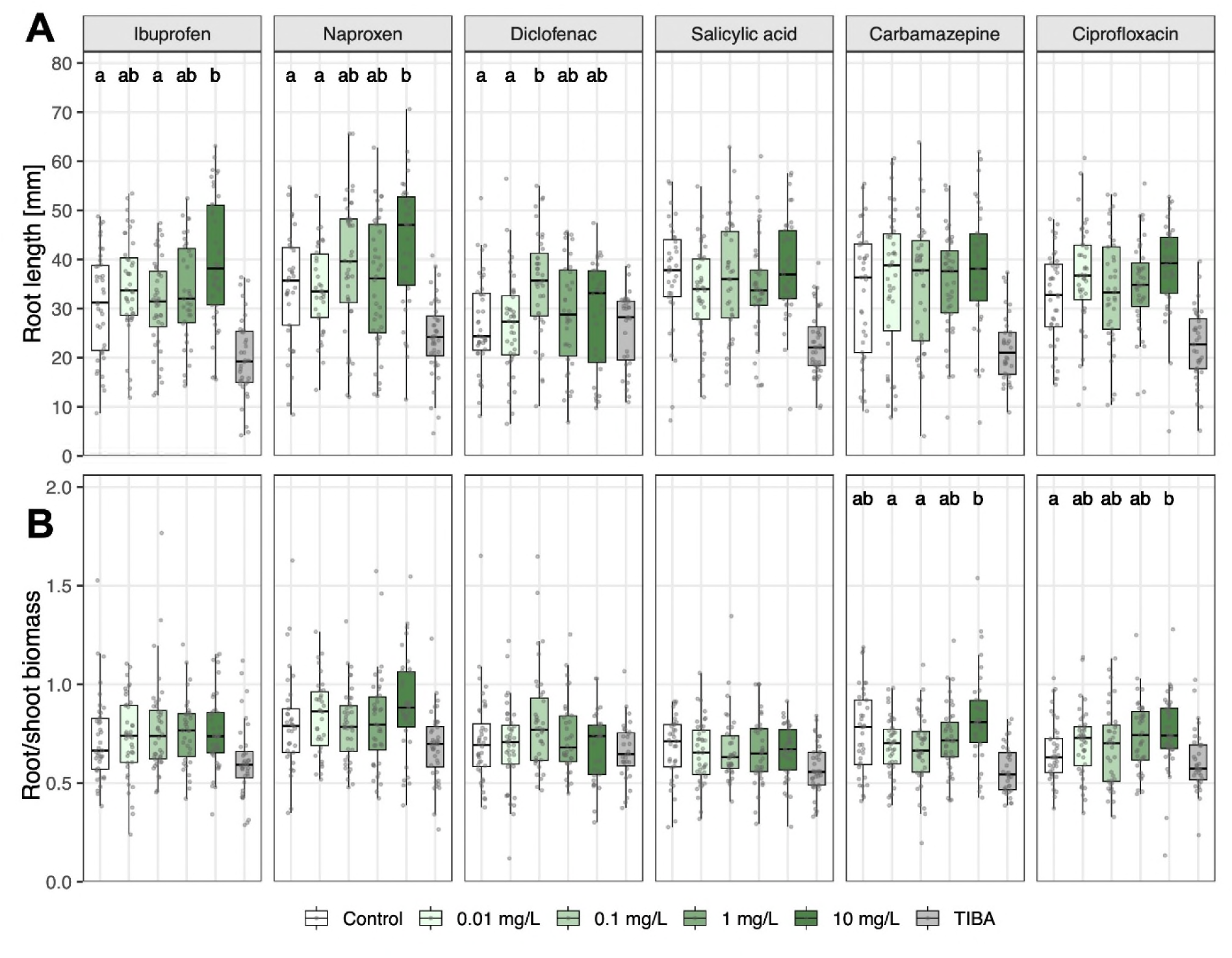
Root length (A) and root-to-shoot biomass ratio (B) of seven-day-old seedlings of *Spinacia oleracea* (spinach) exposed to six pharmaceuticals. Pharmaceuticals were tested in four concentrations, using water as negative control, and 10 mg/L 2,3,5-triiodobenzoid acid (TIBA) as a positive control. Data are presented as boxplots, with medians, interquartile ranges (IQR, boxes), and whiskers extending to the most extreme data points within 1.5×IQR. Individual values are plotted as dots; *n* = 31 - 40 per concentration (from 8 Petri dishes with 10 seeds per dish, 5 germinated seedlings were randomly taken per dish, when available). Different letters indicate statistically significant differences within each pharmaceutical based on Dunn’s post hoc tests following a Kruskal–Wallis test (Bonferroni-Holm-adjusted *p* < 0.05).

In contrast, the root-to-shoot biomass ratio (Fig. 4B) as well as the hypocotyl length (Fig. S4) were only affected by carbamazepine and ciprofloxacin. The root-to-shoot biomass ratio showed significantly higher values at 10 mg/L compared to 0.01 and 0.1 mg/L for carbamazepine and compared to the negative control for ciprofloxacin. The ratios were consistently lower for seedlings exposed to TIBA than for all other treatments (Fig. 4B).

### 3.4. Impacts of pharmaceuticals on lateral root growth of bok choy seedlings

The bok choy seedlings showed clear differences in lateral root development in dependence of the pharmaceutical treatment, varying widely in total lateral root numbers and root length distribution across the three size categories (Fig. 5). In the negative control treatment, roots produced ca. 13 lateral roots, with most roots in C2 (1 - 5 mm), followed by C1 (< 1 mm) and very few in C3 (> 5 mm). That pattern was very similar for seedlings exposed to any concentration of carbamazepine. Under the TIBA treatment, almost no lateral roots were found. When exposed to ciprofloxacin, the strongest reduction in total root numbers was found with increasing concentration, with no lateral roots recorded at 1 and 10 mg/L (Fig. 5).

**Fig. 5.**
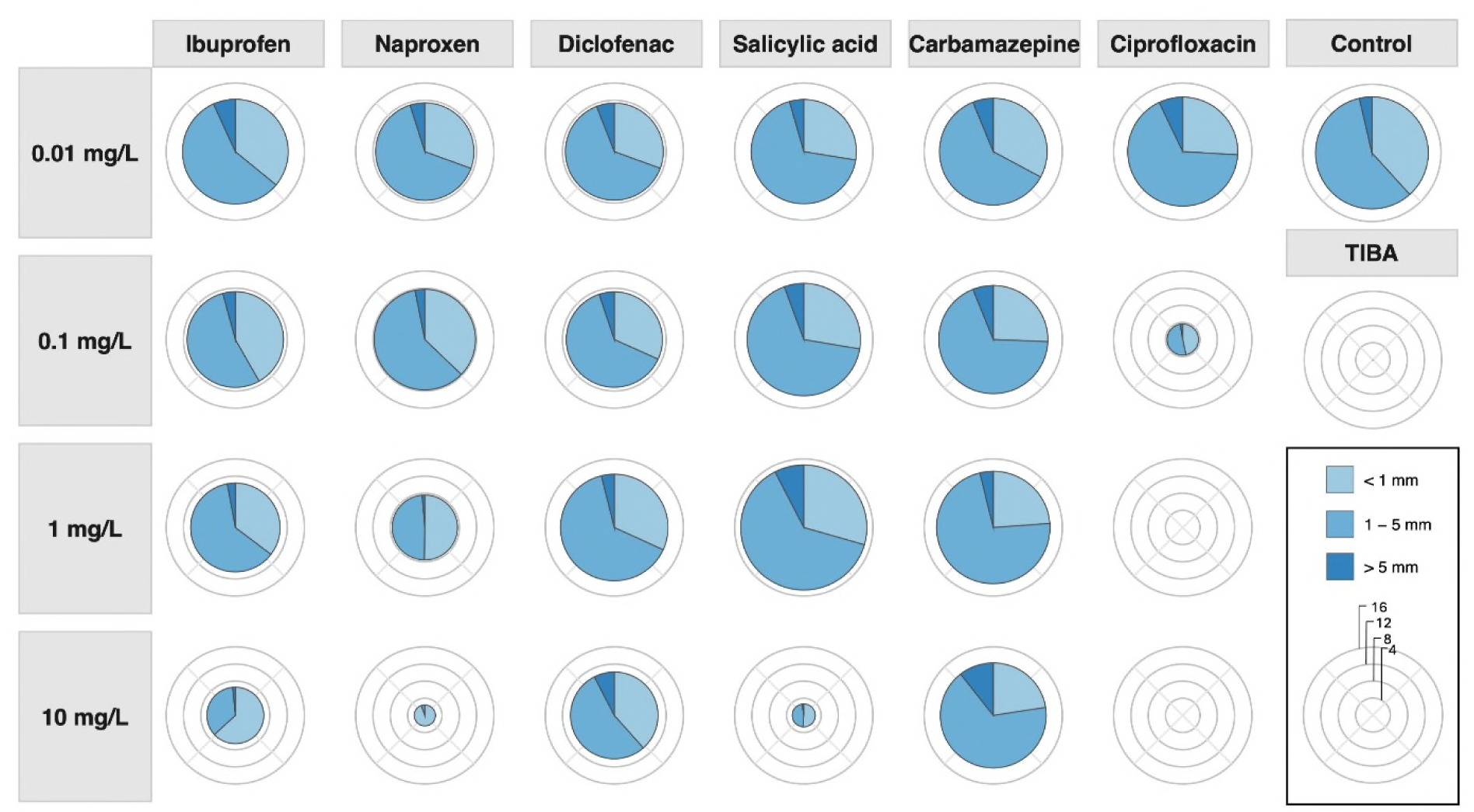
Amount and length distribution of lateral roots of seven-day-old seedlings of *Brassica rapa* subsp. *chinensis* (bok choy) exposed to six pharmaceuticals. Pharmaceuticals were tested in four concentrations, using water as negative control, and 10 mg/L 2,3,5-triiodobenzoid acid (TIBA) as a positive control. Stacked polar bar plots represent the mean number of lateral roots in three length categories for each treatment. The size of each segment indicates the proportion of roots in that category, while the radius reflects the mean total number of roots per treatment; *n* = 29-30.

Also under exposure to the four NSAIDs, concentration-dependent patterns were found. For the treatments with concentrations of 0.01 and 0.1 mg/L, mean total lateral root numbers were mainly slightly lower than those under the negative control treatment, with most roots in C2, some in C1 and few in C3. At the highest concentration, mean total lateral root numbers were substantially lower, especially for seedlings exposed to naproxen and salicylic acid (Fig. 5).

## 4. Discussion

The germination of bok choy and spinach was not affected by any of the six pharmaceuticals at any concentration, in line with our expectation. After seven days, cumulative germination of bok choy and spinach seeds was comparable for nearly all treatments to or even higher than the certified germination rates specified by the supplier (90% for bok choy and 85% for spinach), particularly for bok choy. Our findings are consistent with the results of other studies, which report that the germination rates were unaffected by different pharmaceuticals at similar concentrations (Hillis et al., 2011; Biczak et al., 2024; Kummerová et al., 2024). Even at much higher concentrations, only salicylic acid significantly inhibited the germination of *Lactuca sativa* (1500 - 2000 mg/L), while ibuprofen and diclofenac caused less than 12% inhibition (3000 mg/L) (Pino et al., 2016). Such minimal impact on germination is likely due to the protective function of the seed coat which prevent the pharmaceuticals from reaching the embryo, and to the nutrient reserves within the seed, that allow the embryo to emerge successfully until the primary root develops and comes into contact with the pharmaceuticals (Hillis et al., 2011; Pan and Chu, 2016). In our study, even spinach seeds soaked for 24 hours in pharmaceutical solutions showed no differences in germination rates, indicating that these protective functions last for at least one day. Overall, the robustness of germination appears to be largely independent of seed coat structure and plant family, as revealed by our study and across diverse species from several different families (Hillis et al., 2011; Pan and Chu, 2016).

While the germination remained unaffected, pharmaceuticals had different effects on various early developmental parameters in both bok choy and spinach, indicating that sensitivity to pharmaceutical exposure begins after germination. Impacts were observed especially for bok choy seedlings, which showed an enhanced investment in shoots, while root growth was reduced under exposure to the highest concentration (10 mg/L) of all pharmaceuticals except carbamazepine. This differential investment in shoots versus roots may be explained by impacts of the pharmaceuticals, especially the NSAIDs, on auxin. At least in *Arabidopsis thaliana* it has been shown that ibuprofen, naproxen, diclofenac, and salicylic acid influence auxin-related processes, although through different underlying mechanisms (Tan et al., 2020; Ke et al., 2021; Xia et al., 2023; Cho and Kim, 2021). Ibuprofen disrupts auxin transport indirectly, likely through effects on endocytosis, endomembrane trafficking, and the subcellular localization of auxin transporters (Tan et al., 2020). In contrast, naproxen directly inhibits auxin transport by binding to multiple PIN auxin efflux carriers and inhibiting their transport activity (Xia et al., 2023). Diclofenac has been shown to affect auxin-responsive gene expression, although its precise molecular mechanism within the auxin pathway remains unclear (Cho and Kim, 2021). Salicylic acid, a phytohormone involved in plant immunity and hormonal pathways, is associated with indirect effects on the auxin system by reducing PIN2 mobility at the plasma membrane and its endocytic cycling (Ke et al., 2021; Spoel and Dong, 2024). As auxin is a key regulator of plant development and is primarily synthesized in the shoot apical meristem and young leaves, disruption of auxin homeostasis caused by a decreased transport reduces auxin accumulation in the roots (Olatunji et al., 2017). This may result in reduced auxin signaling in the root meristem, leading to decreased cell division and disrupted lateral root initiation, and thus shorter primary roots and fewer lateral roots. A disrupted auxin gradient may also result in increased auxin accumulation in the hypocotyl and leaves, which can promote cell elongation and thereby contribute to a longer hypocotyl and larger cotyledons (Olatunji et al., 2017). Such mechanisms may also explain our findings. Additionally, NSAIDs have been reported to induce oxidative stress responses in plants, which generate reactive oxygen species (ROS) that reduce cell division and impair root growth and in turn can also influence signaling pathways of hormones, such as auxin (Pasternak et al., 2023; Garduño-Jiménez and Carter, 2024). A suppressed root growth has been likewise found in other species such as *Solanum lycopersicum* and *Lactuca sativa* plants exposed to 2mg/L or 10 mg/L of naproxen or diclofenac, respectively, in seven-day-old seedlings (Siemieniuk et al., 2021, Kummerová et al., 2024).

Antibiotics, such as ciprofloxacin, have also been reported to induce oxidative stress in plants, as indicated by increased ROS levels (Riaz et al., 2017; Hu et al., 2021). In addition, ciprofloxacin directly binds and inhibits chloroplast DNA gyrase in *Arabidopsis thaliana*, which in turn causes chlorosis and disrupts chloroplast division at 10 µM (Evans-Roberts et al., 2016). Such impacts on chloroplasts may also explain the yellow-colored cotyledons of bok choy found in seedlings exposed to 10 mg/L ciprofloxacin in the present study.

In contrast to the NSAIDs and ciprofloxacin, bok choy exposed to carbamazepine exhibited no measurable effects on shoots and roots. As a nonionic compound, carbamazepine can cross root cell membranes more readily than ionic compounds, although its uptake and translocation are also influenced by plant physiology, the concentration, and exposure time (Malchi et al., 2014; Leitão et al., 2021).

Carbamazepine is known to accumulate in plant tissues over time, for example, in leaves of bok choy or *Lactuca sativa*, without causing immediate visible phytotoxic effects ( Li et al., 2018; Stamm et al., 2025). The head biomass of hydroponically grown *Lactuca sativa* begins to decrease at a concentration of 41 mg/L only starting in the fourth week, while root biomass was reduced only at 83 mg/L (Stamm et al., 2025). Thus, although carbamazepine is taken up by plants, it has generally little effects, at least short-term. Overall, the results for bok choy show that different pharmaceuticals can act differently on early development, supporting our hypothesis.

As for bok choy, pharmaceutical treatments had also different impacts on spinach, but overall effects were much less pronounced and partly contrasting, leading namely to a slight root length increase under ibuprofen, naproxen, and diclofenac exposure, and a significant increase in root-to-shoot ratio under carbamazepine and ciprofloxacin exposure. The species-specific results found in our study may be explained by differences in seed size, germination timing, and potential differences in uptake, but also by different experimental treatments. Bok choy has much smaller seeds than spinach and almost all seeds germinated within the first 24 hours, allowing early contact with the pharmaceutical solutions. In contrast, the larger spinach seeds, likely having more nutrient reserves, germinated mainly between 24 and 72 hours. Thus, their primary roots were exposed for a shorter time to the pharmaceuticals. Moreover, spinach was only exposed to one tenth of the solution volume compared to pak choy within the first 72 h, as preliminary assays had shown that spinach seeds did not germinate in general when exposed to a larger volume.

Recent studies have shown that different pharmaceuticals, such as antibiotics, can induce highly species-specific and compound-dependent reactions across plants, ranging from stimulation of root growth to strong inhibition of root elongation and biomass accumulation (Cortés-Lagunes et al., 2026). Effects of NSAIDs were found to depend on the growth stage, sometimes only appearing after several weeks, and leading sometimes to opposite effects on plant development depending on the species. For instance, exposure to 1 mg/L diclofenac enhanced cotyledon opening in *Lactuca sativa* but suppressed it in *Raphanus sativus*, while 1 mg/L ibuprofen promoted primary root development in *Lactuca sativa* but not in *Raphanus sativus* (Schmidt and Redshaw, 2015). Differences may also arise from species-specific sensitivities in metabolic and stress responses, variation in biochemical signaling pathways such as ROS and antioxidant mechanisms (Imani Asl et al., 2024), and genetic architecture, as illustrated by auxin-related genes ranging from 24 in spinach to 119 in *Brassica napus* (Cortés-Lagunes et al., 2026). Although some pharmaceuticals showed no impacts on biomass in our study in the short term, especially in spinach, pharmaceuticals influence plant phytohormones, metabolism, and nutrient uptake at the cellular level (Carter et al., 2015), which may lead to sublethal effects and potential changes in growth over the long term. Further studies are needed to investigate such long-time effects.

## 5. Conclusion

This study showed that pharmaceuticals from different classes did not affect the germination of bok choy and spinach, while the effects on early seedling development were species-specific. In bok choy, higher concentrations of NSAIDs and ciprofloxacin increased shoot and cotyledon growth while reducing root growth and lateral root formation, suggesting disruptions in auxin transport and signaling. Spinach showed only minor and partly contrasting effects. Our findings indicate that sensitivity to pharmaceuticals arises after germination and depends on both the chemical properties of the pharmaceuticals and the plant species. With increasing accumulation of pharmaceuticals in our environment, more severe impacts on plant growth may be expected, since effects are concentration dependent.

## CRediT authorship contribution statement

**Lukas Brokate:** Conceptualization, Data curation, Formal analysis, Investigation, Methodology, Visualization, Writing – original draft. **Caroline Müller:** Conceptualization, Project administration, Resources, Supervision, Validation, Writing – review and editing.

## Acknowledgement

We thank Olajumoke Iqmat Lasisi and Julius Schackmann for help in the germination assays as well as Iman Ayech for help in the lateral roots assay.

## Appendix A. Supporting information

Supplementary data associated with this article can be found in the online version at …

## Data availability

Data and code will be made available upon publication *via* GitHub (https://github.com/PharmaLeaf/Germination-assay.git).

