## Supplementary Table and Figures 1-4 for "Influence of different pharmaceuticals on the germination and early development of two leafy vegetable species"

**Table S1:** pH-values measured for the solutions of the different pharmaceuticals, concentrations, and the controls.

| | 0.01 mg/L | 0.1 mg/L | 1 mg/L | 10 mg/L |
|---|---|---|---|---|
| Ibuprofen | 5.65 | 5.69 | 5.75 | 5.64 |
| Naproxen | 5.93 | 5.75 | 5.63 | 6.05 |
| Diclofenac | 5.58 | 5.68 | 5.68 | 5.75 |
| Salicylic acid | 5.85 | 5.72 | 5.73 | 5.74 |
| Carbamazepine | 5.72 | 5.82 | 5.86 | 5.60 |
| Ciprofloxacin | 6.03 | 6.01 | 6.17 | 6.39 |
| Millipore water | 5.97 | | | |
| TIBA | 4.58 | | | |

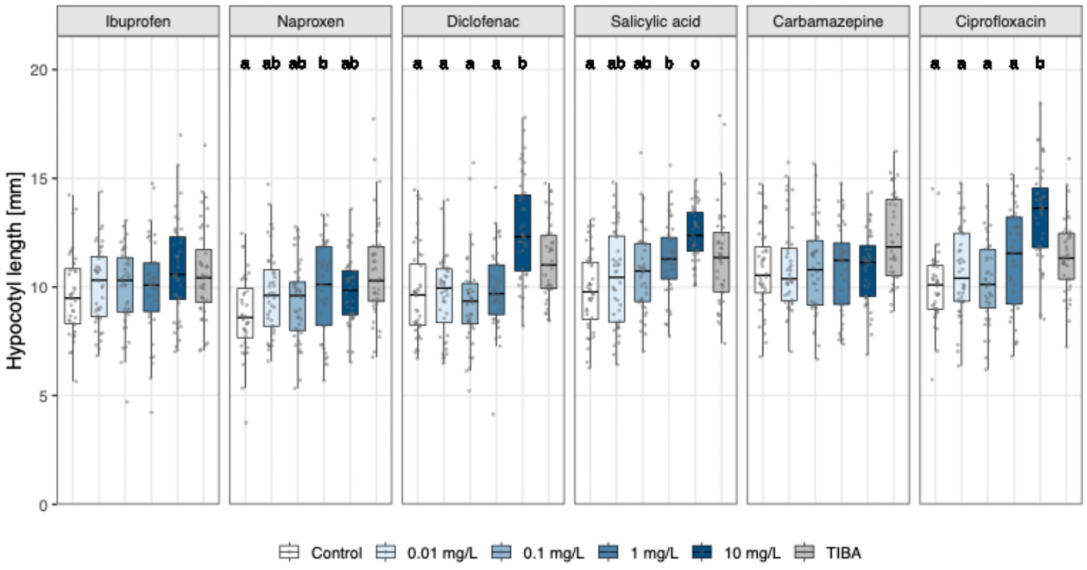


**Fig. S1.** Hypocotyl length of seven-day-old seedlings of *Brassica rapa* subsp. *chinensis* (bok choy) exposed to six pharmaceuticals. Pharmaceuticals were tested in four concentrations, using water as negative control, and 2,3,5-triiodobenzoid acid (TIBA) as a positive control. Data are presented as boxplots, with medians, interquartile ranges (IQR, boxes), and whiskers extending to the most extreme data points within 1.5×IQR. Individual values are plotted as dots; $n = 40$ per concentration (from 8 Petri dishes with 10 seeds per dish, 5 germinated seedlings were randomly taken per dish). Different letters indicate statistically significant differences within each pharmaceutical based on Dunn's post hoc tests following a Kruskal–Wallis test (Bonferroni-Holm-adjusted $p < 0.05$).

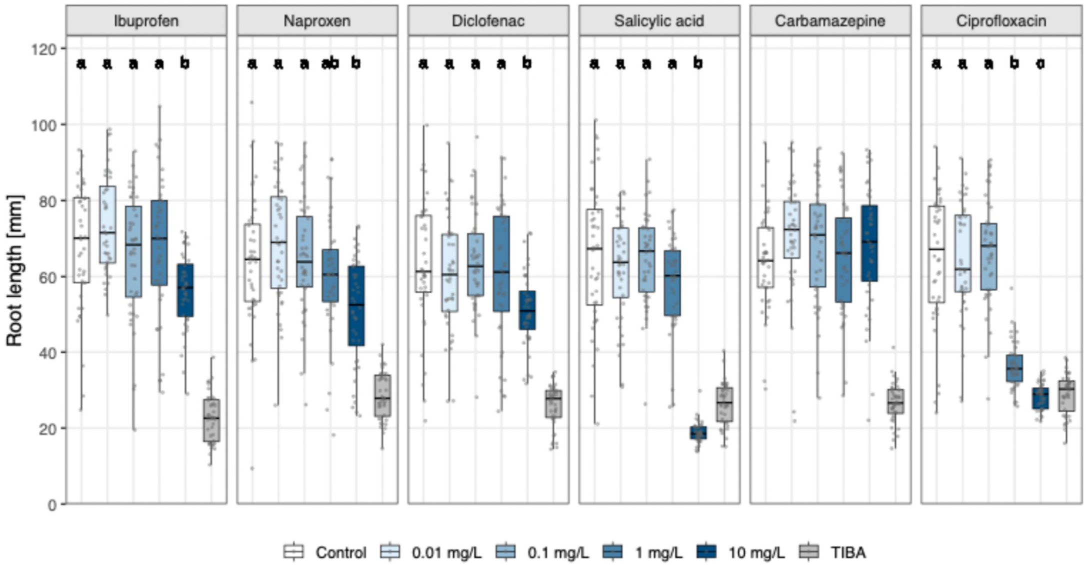


**Fig. S2.** Root length of seven-day-old seedlings of *Brassica rapa* subsp. *chinensis* (bok choy) exposed to six pharmaceuticals. Pharmaceuticals were tested in four concentrations, using water as negative control, and 2,3,5-triiodobenzoid acid (TIBA) as a positive control. Data are presented as boxplots, with medians, interquartile ranges (IQR, boxes), and whiskers extending to the most extreme data points within 1.5×IQR. Individual values are plotted as dots; $n$ = 40 per concentration (from 8 Petri dishes with 10 seeds per dish, 5 germinated seedlings were randomly taken per dish). Different letters indicate statistically significant differences within each pharmaceutical based on Dunn's post hoc tests following a Kruskal–Wallis test (Bonferroni-Holm-adjusted $p < 0.05$).

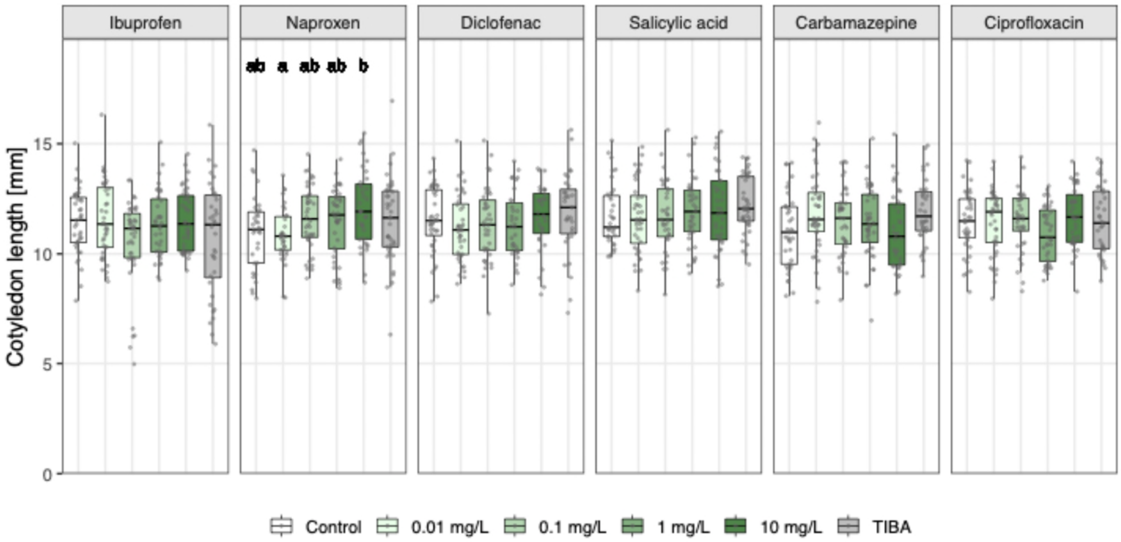


**Fig. S3.** Cotyledon length of seven-day-old seedlings of *Spinacia oleracea* (spinach) exposed to six pharmaceuticals. Pharmaceuticals were tested in four concentrations, using water as negative control, and 2,3,5-triiodobenzoid acid (TIBA) as a positive control. Data are presented as boxplots, with medians, interquartile ranges (IQR, boxes), and whiskers extending to the most extreme data points within 1.5×IQR. Individual values are plotted as dots; $n$ = 31 - 40 per concentration (from 8 Petri dishes with 10 seeds per dish, 5 germinated seedlings were randomly taken per dish, when available). Different letters indicate statistically significant differences within each pharmaceutical based on Dunn's post hoc tests following a Kruskal–Wallis test (Bonferroni-Holm-adjusted $p < 0.05$).

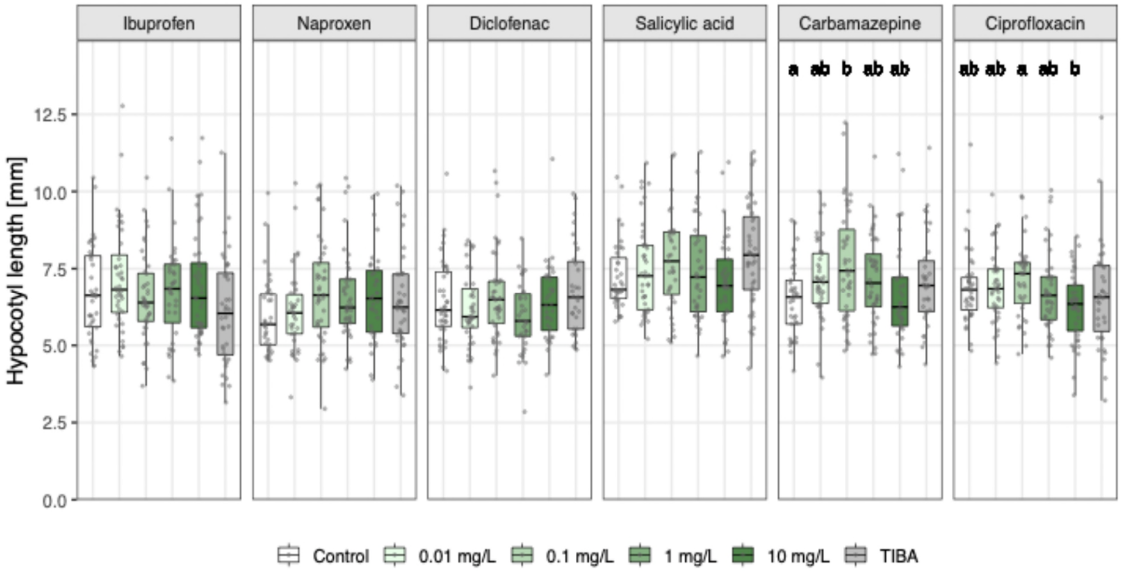


**Fig. S4.** Hypocotyl length of seven-day-old seedlings of *Spinacia oleracea* (spinach) exposed to six pharmaceuticals. Pharmaceuticals were tested in four concentrations, using water as negative control, and 2,3,5-triiodobenzoid acid (TIBA) as a positive control. Data are presented as boxplots, with medians, interquartile ranges (IQR, boxes), and whiskers extending to the most extreme data points within 1.5×IQR. Individual values are plotted as dots; $n$ = 31 - 40 per concentration (from 8 Petri dishes with 10 seeds per dish, 5 germinated seedlings were randomly taken per dish, when available). Different letters indicate statistically significant differences within each pharmaceutical based on Dunn's post hoc tests following a Kruskal–Wallis test (Bonferroni-Holm-adjusted $p < 0.05$).
